# Drought and Productivity Trends in Freshwater Flooded Forest Swamps in the North American Coastal Plain

**DOI:** 10.64898/2026.09.16.752089

**Authors:** Isabel Holland, Kayla Stan, Clay Tucker

## Abstract

Climate change has resulted in an increase in drought conditions within the southeastern United States, impacting a variety of ecosystems and species, including the freshwater flooded forest swamps of the North American Coastal Plain (NACP). With the threat of drought leading to a decline in wetlands and shifts in seasonality patterns in the NACP, there is significant concern about the effects of a drying climate on the net primary productivity of bald cypress-tupelo swamps and the potential for ecosystem degradation or conversion. This study seeks to understand the relationship between the net primary productivity of wetlands dominated by bald cypress-tupelo swamps and drought conditions. We used Moderate Resolution Imaging Spectroradiometer (MODIS) imagery from 2000 to 2024, taken from the 500 largest flooded forest swamps in the southeastern USA using Google Earth Engine, to track the changes in productivity of these swamps over time. The trends in MODIS Fraction of Absorbed Photosynthetically Active Radiation (FPAR), Normalized Difference Vegetation Index (NDVI), and Enhanced Vegetation Index (EVI) were compared against three common drought indices using seasonal decomposition and linear regression along with local bivariate spatial associations. We found that while there were small global correlations between drought and productivity metrics, the impact of droughts on these swamps is highly spatially variable, with specific geographical regions displaying differing impacts of drought and precipitation levels on the vegetation productivity. Coastal and southern regions, including the states of Texas, Florida, and Louisiana, are experiencing a decrease in precipitation but have mixed productivity trends. Meanwhile, more inland and coastal northern areas display an increase in precipitation and wetness. Overall, our results suggest that cypress-tupelo swamp productivity varies based on a variety of factors, including, but not limited to, precipitation and drought levels, but some areas may be at risk of further degradation due to drought under future climate change. Further research is encouraged to fully understand vegetation productivity in relation to environmental conditions in individual cypress-tupelo swamps.

## Introduction

Freshwater forest wetlands in the North American Coastal Plain (NACP) support high levels of biodiversity and provide critical habitats for foraging species and ecosystem engineers (Jorge et al. 2021; Yoezer et al. 2026). These forests are also threatened by multiple sources, including land cover change, climate change, sea level rise, and hydrological alteration (Lang et al. 2024; Yoezer et al. 2026). Since the late 1990s, more than 19,000 km² of forested wetlands in the NACP have been converted to other habitats, primarily scrub-shrubland and marsh wetlands (White Jr et al. 2021). Beyond complete land conversion, an additional 42% of habitats in the NACP are considered highly altered (Noss et al. 2014). Much of the region has been losing carbon through landscape alteration or degradation at an increasingly rapid rate, leading to an increasingly vulnerable region, with potential negative impacts on this highly biodiverse area (Noss et al. 2014; Stan et al. 2026).

Cypress-tupelo swamps are one of the primary freshwater forested wetland ecosystems within the NACP and have experienced a significant decrease in water levels and increased drought conditions over the past half-century due to changes in climate and rapid development (Clem 2019). While mature bald cypress are more abundant in continually flooded areas, seasonal dry periods are required for seeds to properly germinate (Wells 1942). An increase in tupelo and hardwood species has been observed in areas where bald cypress cannot regenerate, leading to a decline in bald cypress-dominated swamps as environmental changes limit their growth (Virginia Department of Conservation and Recreation 2021). Recent work has also found that regeneration in many of these swamps will greatly depend on management practices that reduce salinity levels and allow seedlings to thrive, which is a difficult prospect during rapid sea-level rise occurring in near-coastal swamps (Faulkner et al. 2009; Devereux et al. 2025; Gerdes 2026). With increasing drought conditions lowering the levels of freshwater within cypress-tupelo swamps, there is significant concern about the ability of mature trees to germinate, which impacts tree growth patterns (Doyle et al. 2007).

Dendrochronology has been a successful measure of how long-term changes in both drought conditions and precipitation levels have impacted bald cypress populations over time (Stahle et al. 1985; Stahle et al. 1992; Stahle et al. 2012). Recent tree-ring analysis links bald cypress growth to floodwaters and hydroclimatic change (Copenheaver et al. 2017; Therrell et al. 2020; Tucker et al. 2022); however, there has been limited research into the impacts of recent and regional droughts on the southeastern freshwater flooded forest swamps, the potential impact of drought on vegetation productivity, and the potential for adequate threatened wildlife habitat. The strong response of inland forests to drought-to-pluvial shifts is well understood (Maxwell et al. 2012; Knapp et al. 2016), but research on freshwater-flooded forests globally remains limited, with a need to improve the understanding of the drivers of degradation and ecosystem responses to changing conditions (Yoezer et al. 2026).

With advances in remote sensing technology and computing capacity, there has been an increase in geospatial analysis of flooded forest changes; however, field-based studies, such as dendrochronology, remain dominant in the literature (Yoezer et al. 2026). Those empirical studies are often limited in geographic or temporal scale, which can make it more difficult to parse the relationships between ecosystem function and climate. Historically, remote sensing has had lower-resolution data, and there has been difficulty in harmonizing the large quantities of data pulled from various sensors; however, the rise of freely available cloud-computing geospatial analysis platforms, such as Google Earth Engine, has allowed for the expansion of larger-scale spatiotemporal analyses across fields in agriculture, risk and disaster management, marine science, and environmental change analysis (Amani et al. 2020; He et al. 2018; Liu et al. 2018; Velastegui-Montoya et al. 2023). Much Google Earth Engine research has focused on vegetation and land use and cover changes; however, in more recent years, there has also been an expansion of climate change research that has used these big data approaches (Amani et al. 2020; Velastegui-Montoya et al. 2023; Zhao et al. 2021). These remote sensing products, combined with geospatial methods, enable the collection of environmental and phenological data across broad spatial ranges, temporal scales, and multiple environmental conditions (Khanal et al. 2020).

Therefore, this study aims to fill the gaps in understanding the degradation drivers of the freshwater flooded forest wetlands of the NACP by using remotely sensed data taken from Google Earth Engine to examine the relationship between vegetation indices and drought metrics. Here, we used MODIS vegetation indices (NDVI, EVI, and LAI) from 2000 to 2024 and compared them against the Palmer Drought Severity Index (PDSI) and Standardized Precipitation Index (SPI) taken from GRIDMET and precipitation from CHIRPS across the same 2000 to 2024 period. These time series were decomposed using Seasonal-Trend decomposition using LOESS (STL), and the trend component was assessed for the direction and magnitude of change using linear regression. The indices were then correlated and assessed for spatial correlations using Lee’s L Spatial Association Statistic.

## Methodology

### Study Area and Site Selection

The NACP is located in the southeastern United States, stretching from eastern Texas to the Atlantic coast of North Carolina, and is characterized by flat, low-lying terrain (Lang et al. 2024; Stroud 2025). This coastal plain is home to a variety of plant communities, including bald cypress-tupelo swamps, maritime and estuarine wetlands, and longleaf pine (*Pinus palustris* Mill.) savanna, all of which contribute to a comparatively large area of freshwater-flooded forests in the United States. As one of North America’s largest biodiversity hotspots, the NACP and its wetlands are home to a wide variety of vascular plants, fish, and reptile species, making them a critical area to study for changing pressures (Goebel et al. 2001).

We selected the locations of the 500 largest freshwater-flooded forest swamps in the NACP based on data from the National Oceanic and Atmospheric Administration’s C-CAP High-Resolution Land Cover data. Flooded forest swamps were identified using the NOAA Coastal Change Analysis Program, which produces regional land cover data every five years from Landsat data. For this dataset, only areas that are likely to have changed are reclassified every five years, and for this work, the Palustrine Forest Wetland classification from the 2021 dataset was used. The 2021 map has a resolution of 30 m and an overall accuracy of 85%; however, it is considered to have a more reliable cover identification at a 60 m resolution. The Palustrine Forest Wetland class, in particular, has a user accuracy of 91.6% (National Oceanic and Atmospheric Administration 2025; McCombs et al. 2016).

The 500 largest Palustrine Forest Wetland forest areas were each randomly sampled five times, leading to a total of 2,500 points being assessed. All flooded forest areas were greater than 21 km², with an average swamp area of 58 km² and the largest wetland size of 1,062 km². Each point was a minimum of 500 m apart from other points so that no MODIS pixel was sampled more than once. Additionally, all points had at least 25% of the 500 m pixels composed of flooded forest land cover, and 90% of the points were classified as >50% flooded forest. These thresholds were used to reduce the impact of edge effects and habitat fragmentation on the analysis. Each flooded forest sample site was treated independently. Although these data could be aggregated by the wetland, because microtopographical and hydrologic variation within forested wetlands in the coastal plain has been found to impact carbon dynamics at a seasonal level, each sample was treated as a separate site (Miao et al. 2017). Given the sampling parameters, there may be a small degree of temporal and spatial autocorrelation due to shared environmental conditions within smaller sampled forests.

### Vegetation and Drought Data

The vegetation indices we used included the normalized difference vegetation index (NDVI), enhanced vegetation index (EVI), and leaf area index (LAI), which were generated from MODIS imagery. The normalized difference vegetation index and enhanced vegetation index are proxies for the health and productivity of vegetation using infrared and visible light radiation, with EVI considering corrections for atmospheric noise (Kandasamy et al. 2013). The leaf area index determines the amount of leaf area that can be used for photosynthesis and, therefore, the productivity of vegetative areas. NDVI values range from -1 to +1, with higher values indicating dense, healthy vegetation and lower values indicating bare ground, urban areas, or snow. EVI typically has similar ranges, with higher values indicating denser and healthier vegetation. This index corrects for background and atmospheric noise and does not suffer from saturation that can occur in NDVI. We extracted MODIS 13A1.061 Terra Vegetation Indices 16-Day Global 500 m imagery from 2000 to 2024 using Google Earth Engine to understand the changes in NDVI and EVI flooded forest swamp phenology (Didan 2021). This sensor has a temporal resolution of 16 days and a spatial resolution of 500 m. We used imagery from MODIS MCD15A3H.061 from 2000 to 2024 for sensing LAI vegetation productivity, with level 4 accuracy and a temporal resolution of every four days (Myneni et al. 2015). A level four accuracy represents the highest level of data processing quality provided by MODIS, as well as a high validation level for climate-based studies (NASA 2021).

We collected precipitation data from the Climate Hazards Center InfraRed Precipitation with Station data (CHIRPS) satellite with a daily temporal resolution and a spatial resolution of 0.05° based on longitude and latitude (Funk et al. 2015). Drought data for the standardized precipitation index (SPI) and the Palmer Drought Severity Index (PDSI) were sourced from the 4-km daily Gridded Surface Meteorological (GRIDMET) dataset (Abatzoglu 2012). CHIRPS data measure precipitation levels by combining ground-based measurements with satellite imagery (Funk et al. 2015). The PDSI utilizes regional drought data to measure long-term droughts using soil moisture measured from monthly precipitation and temperature data (Alley 1984). The Standardized Precipitation Index (SPI) is a common drought index that uses monthly precipitation from a set time frame to determine the probability of a particular observed precipitation dataset (McKee et al. 1993). The SPI is particularly useful because it normalizes differences in precipitation across wetter and more arid areas (McKee et al. 1993). While these indices indicate levels of drought, they differ in their calculation complexity, providing unique insights into the potential impact on vegetation productivity. We calculated all drought and precipitation indices from 2000 to 2024 to allow for a comparison with the vegetation indices.

### Analysis

#### Time-Series Decomposition

We extracted all drought indices, CHIRPS precipitation, and MODIS vegetation productivity indices using Google Earth Engine from 2000 to 2024. Gaussian smoothing was applied to the trends for the points for each index. This smoothing technique is used to process data into bell curves to smooth data trends and aid in the reduction of noise when applied to remote sensing data (Kandasamy et al. 2013).

These smoothed time series were then further split using Seasonal-Trend decomposition using LOESS (STL). STL uses locally weighted scatterplot smoothing to separate various trends within the data by capturing nonlinear relationships in comparison to local regressions (Cleveland 1990). This results in the splitting of trends into seasonality, trends, and remaining fluctuation components in the smoothed data (Abbes et al. 2018; Wang et al. 2022). In this study, we only assessed trend data. We then assessed the annual trends from the decomposition using linear regression. Linear regressions were generated for each swamp point for each vegetation and drought index. The directionality and significance of the trend were extracted for each regression.

#### Productivity Hotspot Analysis

To assess the spatial clustering of productivity changes, we tested each index for local hotspot clustering using a Getis-Ord Gi* hotspot analysis using k-nearest neighbors. K-nearest neighbors is preferable to using a fixed band distance given random sampling within wetlands. Getis-Ord Gi* compares the local trends in productivity or drought against the nearest eight points (Getis & Ord 1992). We then calculated the local Gi* value for each point using the following equations:

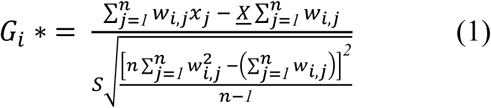

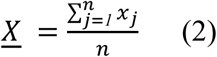

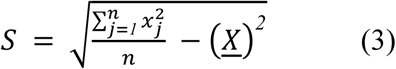

where x_j_ is the local sum for feature j, w_i.j_ is the spatial weight between adjacent features i and j, and n is the total number of features (Yi et al. 2019). The Getis-Ord Gi* statistics were then assessed based on their confidence interval (90%–99%), with negative Gi* representing a decrease in productivity across all productivity metrics. We compared hotspots across all three productivity metrics to determine the similarity of trends across the indices. The similarity between hotspot patterns was calculated using a fuzzy similarity score ranging from 0 to 1, where values approaching 1 indicate greater spatial agreement between hotspot significance confidence intervals (Hagen 2003). The score incorporates both the similarity of significance confidence interval levels (99%, 95%, and 90%) and the spatial agreement of neighboring features. This allows for partial similarity between closely related significance categories or spatially proximate patterns to be accounted for in the score. Global similarity was calculated as the mean similarity across all corresponding features.

#### Spatial Associations Between Productivity and Hydroclimatic Variables

We then correlated productivity trends with three hydrology metrics: precipitation changes, PDSI trends, and SPI trends. To determine local-level relationships between vegetation changes and changes in moisture, we ran spatial associations using Lee’s L statistics. Spatial associations were chosen for this study as they are designed to correlate local-level trends based on the extent to which the data overlap while accounting for spatial homogeneity and lag. Our values ranged between 1 and -1, with positive values indicating a high correlation and similar neighborhood patterns between the two variables, while negative values indicated a negative correlation between the two variables, with high values in one variable and low values in the other. Our data analysis resulted in a statistical measure of clustering, which is an extension of spatial autocorrelation (Getis 2001). The Lee’s L statistic, or bivariate spatial association, measures how related two different sets of spatial data are to one another and is calculated by correlating local averages, smoothing, and applying spatial weights in order to generate an autocorrelation (Lee 2001; Lee 2009). Spatial associations have been found to be especially useful for understanding data that are tied to the underlying causation of trends (Getis 2001). We generated a total of fifteen associations, displaying the spatial patterns and clusters of the impact of drought on vegetation productivity trends.

## Results

### Flooded Forest Productivity Trends

Flooded forests across the NACP displayed mixed productivity trends. An increasing NDVI was recorded at 60% of the points (Figure 1a), whereas the LAI increased at 48% of the points (Figure 1e). In addition, 51% of the points recorded a significantly decreasing LAI over the past 25 years (Figure 1f), whereas 39% of the points had a decreasing NDVI (Figure 1b). The EVI showed similar trends, with 53% of the points recording significant increases (Figure 1c) and 45% recording significant decreases in productivity (Figure 1d). There are relatively stable hotspots and cold spots of change, with 0.84 similarity in the hot and cold spots between NDVI and EVI. LAI had slightly different trends, with 0.77 similarity in hotspots compared to NDVI and 0.76 compared to EVI. Overall, the combined median annual change was -0.002 for NDVI, - 0.023 for LAI, and -0.002 for EVI (SI Figure 1).

**Figure 1.**
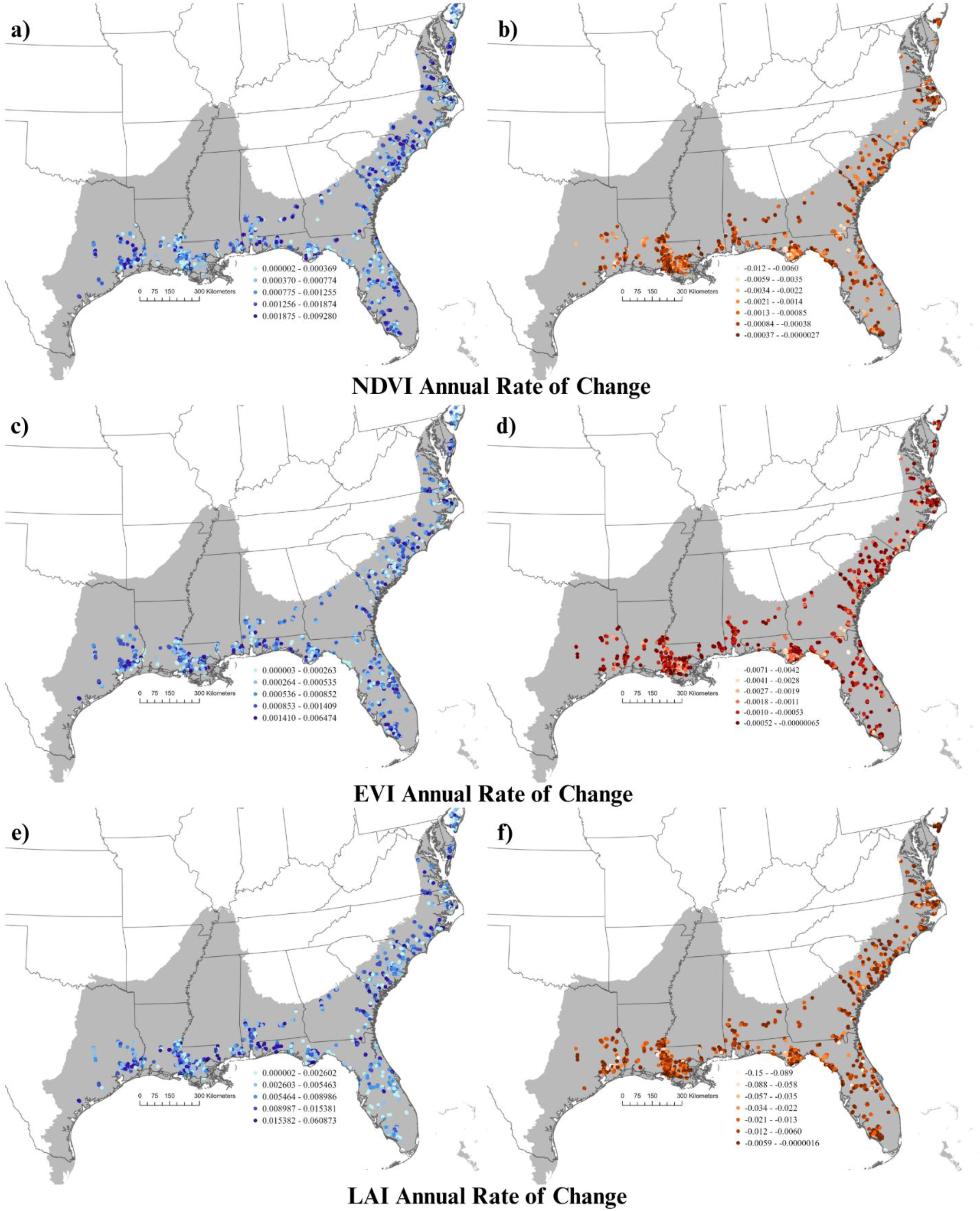
Annual productivity trends of three common indices of the 5 random samples within the 500 largest flooded forest swamps in the North American Coastal Plain. Blue points (left panels) indicate sample points where average productivity is increasing over the past 25 years, while red points (right panels) are areas of productivity decreases. In each case the unit is the Δ productivity index/year. Productivity trends include NDVI (a,b), EVI (c,d) and LAI (e,f).

Vegetation decreases were most notable in the southern coastal regions along the Gulf of Mexico, with clusters of decreasing productivity in Louisiana, the western coast of Florida, and coastal Georgia, South Carolina, and North Carolina (Figure 2a–c). Notable clusters include those in Louisiana along the Mississippi River between St. Catherine Creek and Three Rivers Wildlife Management Area, along Highway 90 between Ellsworth and Morgan City (Bayou Black), near Lake Fausse Pointe, and near the Calcasieu River. Notable clusters in Florida include the Apalachicola River Wildlife and Environment Area near Devil’s Swamp and Otter Creek, the Big Bend Wildlife Management Area, the lower Suwannee Wildlife Refuge, and the Waccasassa Bay Preserve (Figure 2; SI Figures 3–6). In contrast, hotspots of increased productivity are more highly clustered in Texas, the interior flooded forests of Georgia, South Carolina, North Carolina, and further north. Additionally, hotspots were observed in the central parts of Florida (Figure 2). There is also a cluster of increased productivity in Louisiana between East Grand Lake and Bayou Pigeon Road (Figure 2).

**Figure 2:**
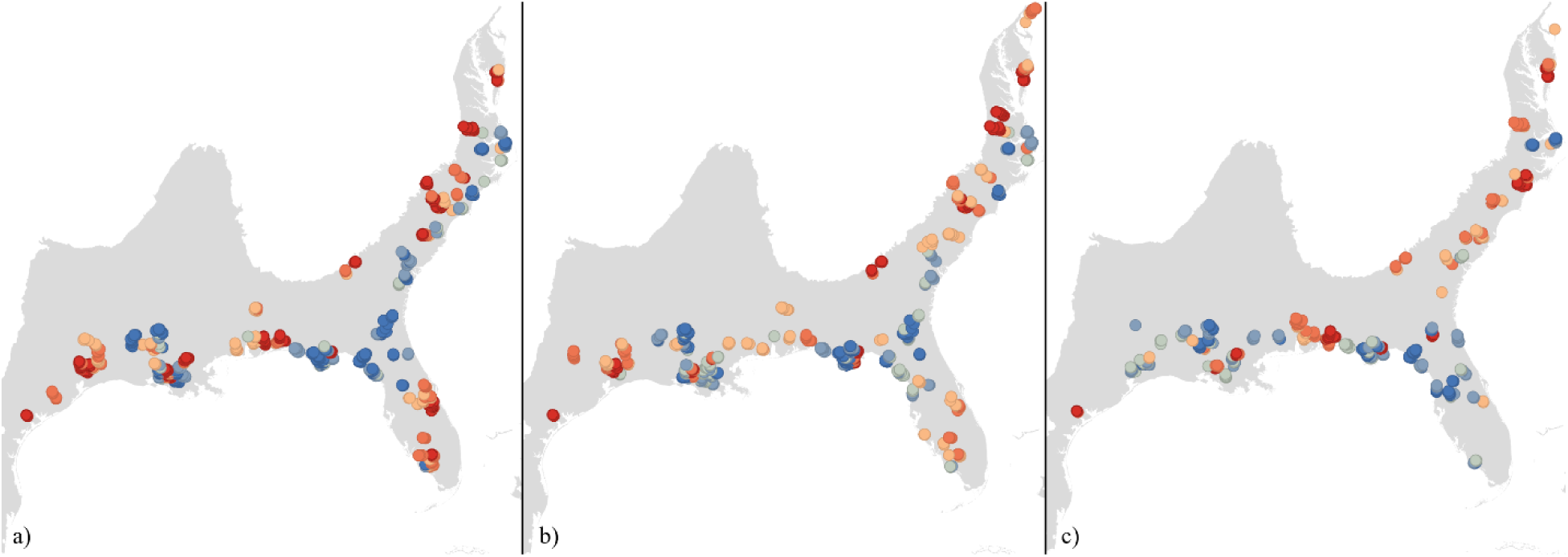
Productivity hotspots of a) EVI, b) NDVI, and c) LAI with red indicating statistical clusters of increasing productivity and blue indicating statistical clusters of decreasing productivity.

**Figure 3.**
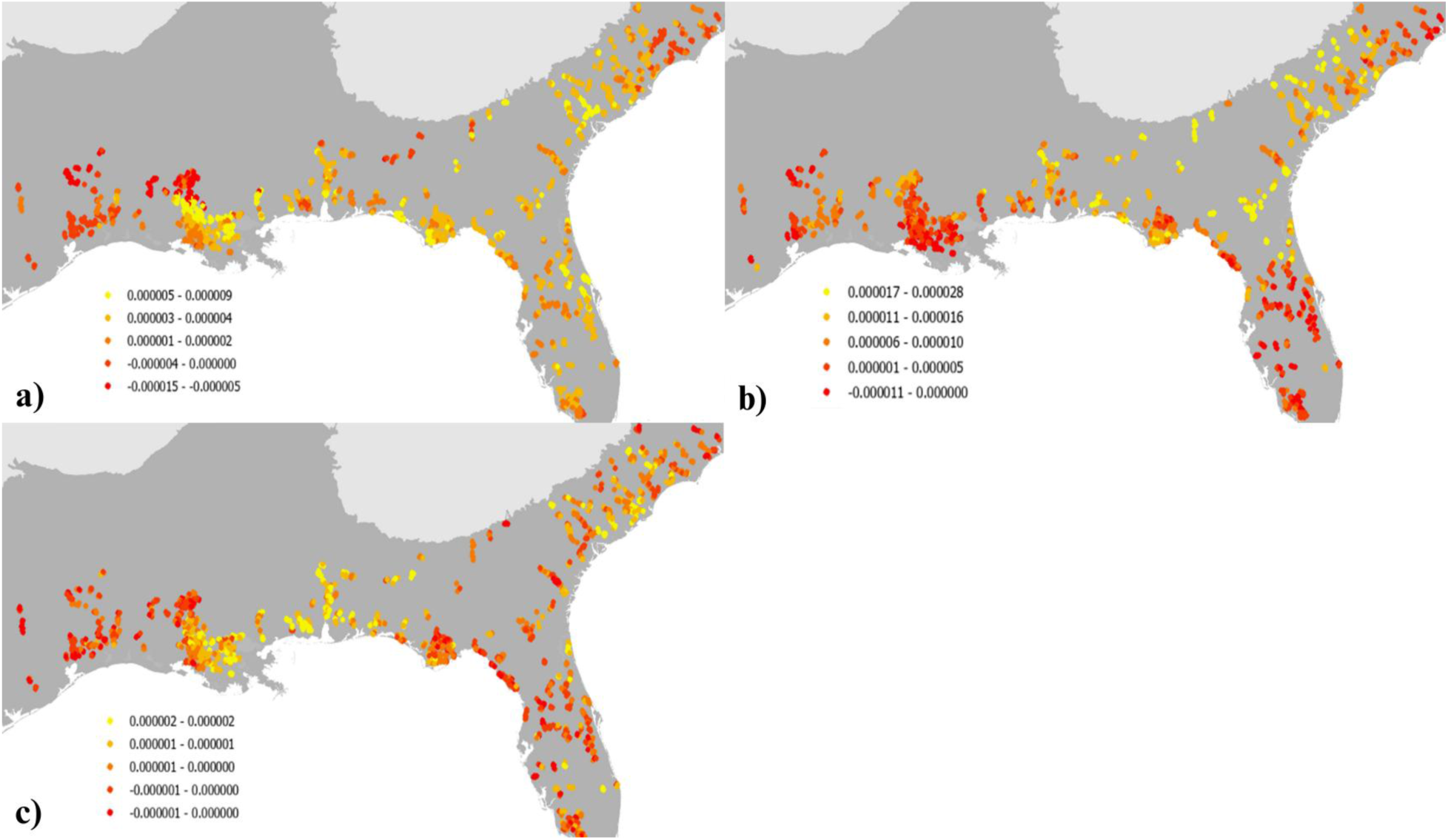
A map displaying the annual average change in precipitation (mm/yr) based on CHIRPS data (a), with red indicated stronger decreases in precipitation. The changes in drought indices include the annual average changes to PDSI (b), and SPI (c). For each index, red points indicate a higher level of drought in the area.

### Precipitation and Drought Condition Trends

Across all three drought indices, most points indicated trends towards wetter conditions across the NACP in the last 25 years. There were increases in PDSI across 84% of points, precipitation across 76% of points, and SPI across 60% of points (Figure 3a–c). The median change in the PDSI was 6.7 × 10⁻⁶, while that in the SPI was 7 × 10⁻⁸. Daily precipitation increased on average by 3 × 10⁻⁶ mm; however, the annual changes across all points were strongly left-skewed (skewness: -1.67, kurtosis: 7.49; SI Figure 2). In general, while the changes in drought and precipitation were significant across many of the points, the trends accounted for very little of the data variance. Precipitation had the lowest average R² of 0.08, whereas PDSI had an average R² of 0.32 (SI Figure 2).

When we consider the spatial clustering of hydrologic changes, only PDSI and precipitation had sufficient clustering to find hotspots of change. Hotspots of increased precipitation included areas in Louisiana between New Orleans and Baton Rouge, along the Mississippi and Pascagoula Rivers, along the eastern coast of Florida, and in many of the flooded forests in South Carolina (Figure 3a; SI Figures 3–7). The clusters of decreasing precipitation were found in North Carolina, much of Texas, and further north in Louisiana. These trends are not consistent with the more complex PDSI; coastal Louisiana is one of the largest clusters of decreasing PDSI, which indicates increased drought. Increased drought clusters are also found in Florida around the Suwannee River basin, throughout the south and central flooded forests of Florida, on the northern side of the Albemarle Sound, and northeast of Houston (Figure 3b; SI Figures 3–7). Increases in PDSI, or decreased drought, can be found in northern Louisiana, along the Pascagoula River, along the coastal Apalachicola management areas, and across the flooded forests from Alabama northeast to the border between North Carolina and South Carolina (Figure 3; SI Figure 3). When the hotspots between precipitation change and PDSI change are compared, there is a lower similarity score compared to the productivity metrics (similarity score: 0.55).

### Productivity and Drought Interaction

When we considered spatiality in the correlations, we found that clustering patterns were present between the trends of drought and vegetation productivity. Twelve percent of flooded forest swamps were experiencing decreasing trends in drought and increased vegetation productivity, with hotspots found in swamps in the inland areas of Alabama, North Carolina, and South Carolina (Figure 4). These findings were heavily reflected in the SPI and PDSI indices (Figure 4a,c,d,f,g,i). Eighteen percent of the flooded forest points displayed trends of increased drought and decreasing trends in vegetation productivity, with hotspots in Louisiana, central Florida, and the Atlantic coast, which were seen across all indices, but primarily with PDSI (Figure 4c,f,i). As this is the largest analyzed percentage, our results show that the greatest number of cypress swamps is experiencing greater levels of drought and negative impacts on vegetation productivity. Sixteen percent of the studied swamps were undergoing decreasing drought trends, as well as decreased levels of net primary productivity, and were found in South Carolina, Georgia, and on the coast of Florida. Swamp points with increasing drought trends and increasing vegetation productivity were located in southern Florida and east Texas. Forty percent of swamp locations experienced non-significant spatial relationships between recorded productivity and drought trends (Figure 4).

**Figure 4.**
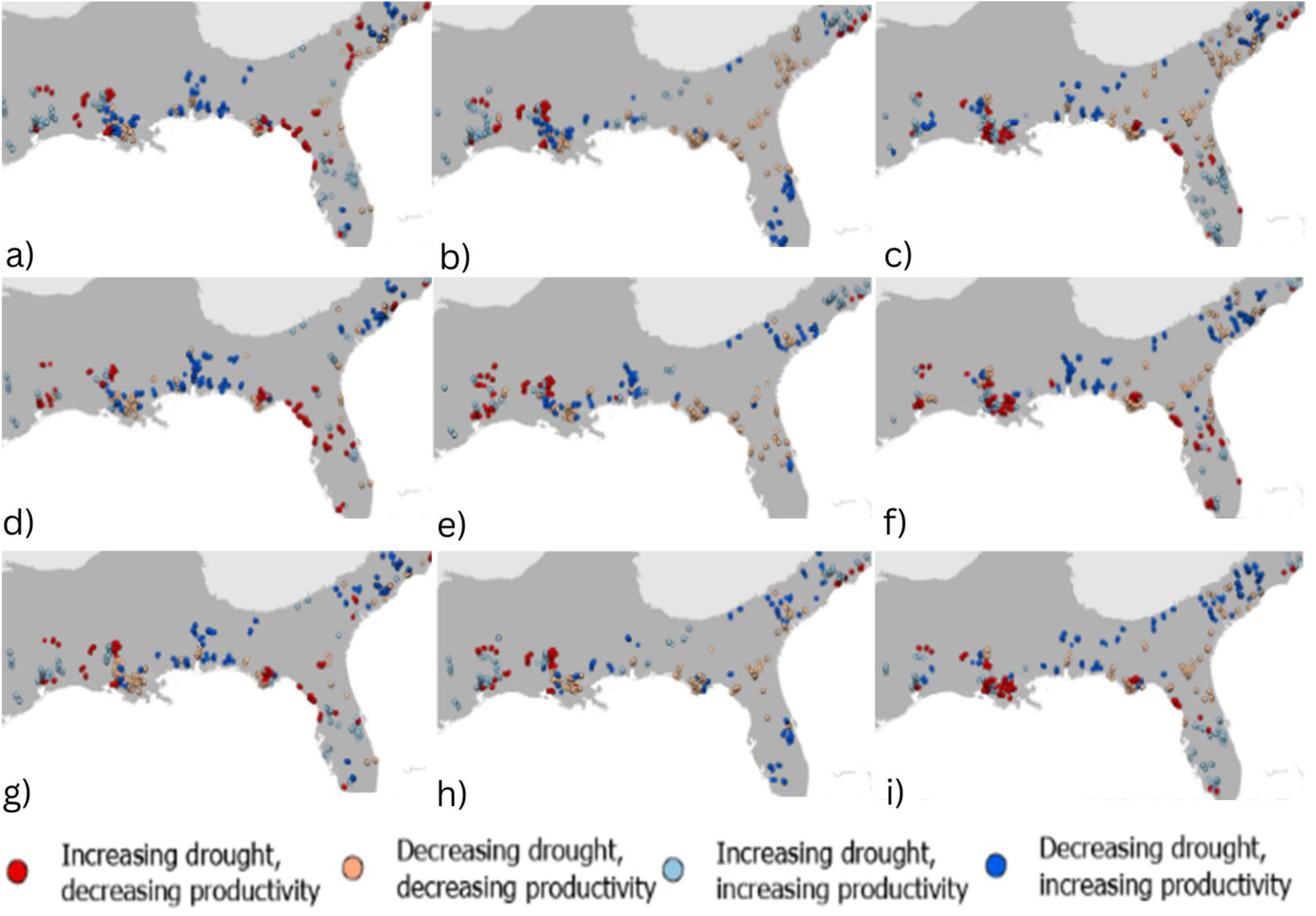
The spatial association between changes in a) SPI and EVI, b) precipitation and EVI, c) PDSI and EVI, d) SPI and LAI, e) precipitation and LAI, f) PDSI and LAI, g) SPI and NDVI, h) precipitation and NDVI, and i) PDSI. Dark red indicates increasing drought and decreasing productivity and dark blue indicates decreasing drought and increasing productivity, or where flooded forests are on trend with changes in drought. Light red indicates decreasing drought and decreasing productivity and light blue indicates increasing drought and increasing productivity, or areas where there may be other influencing factors.

Spatial associations between EVI and the drought indices resulted in spatial patterns of decreased wetness and vegetation in southern Louisiana, the coasts of Florida and Texas, and the coastal areas of North Carolina (Figure 4a–c). Significant decreases in wetness with high vegetation productivity were noted in much of southern Florida, eastern Texas, and central Louisiana. High wetness with low vegetation productivity was recorded on the coasts of South Carolina and Florida. Increases in wetness trends and vegetation productivity were found in swamps in Mississippi, Alabama, and the Carolinas, all of which are not located in coastal areas.

In the spatial associations between NDVI and drought indices, increases in both wetness and vegetation productivity were found within the areas of the inland, northern range of the flooded forest swamps (Figure 4g–i). States including North Carolina and South Carolina in the northern range and Mississippi and Alabama displayed increased vegetation and wetness along the coasts. Areas with increased wetness and lower levels of vegetation productivity include the southern and coastal parts of Louisiana and Florida, as well as protected areas in these states. Lower wetness and increased vegetation productivity trends were noted in eastern Texas, southern and central Florida, central Louisiana, and the inland areas of North Carolina and Georgia. Decreases in wetness and vegetation productivity were observed in the northern areas of Florida and Louisiana.

Spatial associations between LAI and the various drought trends resulted in high amounts of both wetness and vegetation productivity throughout Alabama, Georgia, and South Carolina, with swamps located along the banks of rivers (Figure 4d–f). High wetness and lowered productivity trends were primarily observed in southern Louisiana, on the coast of Florida, and on the coasts of Georgia and South Carolina. Increased drought trends from decreasing wetness and vegetation productivity were found in central Louisiana, east Texas, and central and coastal Florida. Decreases in wetness and increases in vegetation productivity were spatially located in the inland areas of Georgia, the coast of North Carolina, and central Texas.

## Discussion

### Drought Trends and Flooded Forest Productivity

The results of this study indicate that some cypress-tupelo swamps are experiencing significant drought conditions in the southern and coastal areas of the NACP, impacting the productivity of vegetation. Overall, this study found that 18% of the flooded forest swamps are experiencing increasing drought trends and decreasing levels of vegetation productivity, with hotspots located in Louisiana and the Atlantic coast, indicating degradation in these areas. Additionally, while swamps in South Carolina, Georgia, and on the coast of Florida are becoming degraded without increasing drought trends, many cypress swamps in Texas, Louisiana, and southern Florida are correlated with increases in drought. This drought impact is particularly true in southeastern Louisiana, which mirrors patterns found in other studies from the flooded forest swamps in the southeastern USA that suggest that drought conditions lead to increases in saltwater inundation and decreased growth in cypress, while large influxes of water late in the growing season can lead to additional bursts of growth in cypress trees (Therrell et al. 2020; Tucker et al. 2022; Day et al. 2024).

Although the average annual changes in NDVI, EVI, and LAI are low overall, over the course of the last two decades, these have led to decreases in average annual productivity of between 1% and 3%. If these trends continue under further climatic changes, 10% of the swamps could experience more than a 10% drop in average productivity by the end of the 21st century, and 2% of these wetlands would have more than a 25% decrease in productivity. This drop is further magnified if we consider the total overall productivity instead of the average annual vegetation indices. Average NDVI alone has a curvilinear relationship with gross primary productivity in the United States, and flooded forests typically have higher vegetation indices than more northern points, making these drops in productivity likely more pronounced in gross primary productivity and biomass metrics (Phillip et al. 2008).

### Factors Influencing Flooded Forest Swamp Degradation

Overall, our work indicates that there are other factors beyond drought, at regional and local levels, that impact the health and productivity of these ecosystems, which is supported by previous research. Historical dredging, logging, and drainage of prime bald cypress-tupelo swamp ecosystems have negatively impacted ecosystem services and the distribution of native species, as well as limited tree growth (Stahle et al. 2012). Current management practices to mitigate flood discharge in the Atchafalaya River Basin, Louisiana, have been suggested to be inadequate (Kroes et al. 2022). Furthermore, isolated wetlands in the NACP are increasing as the demand for infrastructure and agriculture continues. Frameworks for measuring these impacts have found that more ecological harm occurs in wetlands surrounded by agriculture (Stuber et al. 2016). In coastal areas, cypress-tupelo swamps are facing the growing threat of increased tidal flooding as a result of eustatic and relative sea-level rise. Areas with constant flooding resulted in decreased overall bald cypress growth compared to simulated periodic flooding. Saplings were significantly impacted, as seasonal dry periods allowed germination (Megonigal and Day 1992). Cypress-tupelo swamps are likely to experience an increase in extreme weather events, such as hurricanes, that impact the productivity of these ecosystems. However, hurricane impacts on NACP ecosystems are also nuanced. Chronically stressed environments experience growth limitations from multiple factors (Tucker et al. 2025), and individual species have differential reactions to the associated wind, precipitation, and storm surge damage (Tucker et al. 2018; Mitchell et al. 2019; Rutledge et al. 2021; Tucker et al. 2025). Bald cypress-dominated swamps have also been found to experience relatively quick regeneration after extreme hurricane damage compared to hardwoods in the same locations (Ramsey et al. 1998).

Recent studies have found that salinity levels in bald cypress swamps are increasing yet have not been recorded to have a significant impact on the overall growth patterns of mature trees (Doyle et al. 2007). Seedlings cannot germinate in water with high salinity levels, and forest die-off from salinity levels that exceed 2 ppt cannot be regenerated by management practices (Day et al. 2024; Faulkner et al. 2009). Inland trees typically display a lower tolerance to high salinity levels, with coastal trees being more adapted to increasing salinity levels owing to regular periodic flooding and consistent precipitation levels that allow for salinity discharge (Day et al. 2024). Precipitation is the main driver of reduced salinity levels in both coastal and inland cypress-tupelo swamps. As drought trends threaten these ecosystems, there is concern over the availability of precipitation to reduce salinity levels and prevent the further spread of these “ghost forests.”

### Implications for Biodiversity and Wildlife Habitat

As bald cypress swamps experience varying degrees of drought impact, there is concern regarding the impact of decreased wetness on the animal species that inhabit and roost within these swamps. With more extensive drought patterns, there will likely be limitations on insect populations and the availability of quality bat roost and foraging spaces (Clement and Castleberry 2013). More broadly, changing conditions and increases in drought have been found to decrease the diversity in wetland ecosystems, particularly impacting species endemic to these environments (Virginia Department of Conservation and Recreation 2021; Thorne et al. 2016). With increases in soil disturbances, there has been a shift in seed recruitment towards fugitive species, which can rapidly colonize an unstable area. Less frequent flooding in an area has also been found to shift wetlands in the Southeast towards a more terrestrial species composition; however, these systems are thought to have the highest species richness when there is a balance of disturbances from events such as droughts, allowing successional stage progression (Kirkman 1995).

### Limitations and Future Research

There are limitations to this type of index analysis that should be considered. Using a singular sensor is not always adequate for the collection of phenological data and could have introduced some bias in this study, as only MODIS data were utilized for the phenology of cypress-tupelo swamps (Zou et al. 2023). Abbes et al. (2018) tested three different decomposition methods for vegetation indices and assessed phenological changes, with BFAST and MRA-WT being the most accurate. Decomposition was a prominent form of analysis in this study, allowing for the determination of vegetation and drought trends. MODIS imagery has some drawbacks when measuring vegetation. When evaluating tree cover, trees of lower heights are often unsensed and exempted from data, leading to potential miscalculations of vegetation indices (Yang and Crews 2019). CHIRPS data have also been found to not fully capture extreme event data, particularly under hurricane conditions (Vega-Camarena et al. 2026), which may lead to bias in the precipitation relationship. CHIRPS has been found to be most accurate at annual scales and is useful for assessing seasonal droughts; however, it has been found to underestimate precipitation, which should be taken into account (Alisilibe et al. 2023).

Previous studies have found inaccuracies when assessing seasonal changes caused by extreme weather events, including severe drought conditions, which is a concern for the study areas in this project (Zou et al. 2023). Future work could utilize sub-pixel models, which allow for the understanding of small-scale coastal changes caused by ecosystem changes, with a particular emphasis on the time scale used in imagery processing (Li and Gong 2016). Integrating vegetation indices over the entire growing season has also been more closely linked to overall site productivity, which has improved plant ecosystem monitoring and could be pursued in the future (Yan et al. 2022). Monitoring plant community diversity can be used as an indicator of ecosystem health in wetlands, as well as to understand shifts in composition caused by a variety of climate and environmental factors (Dronova and Taddeo 2022). LIDAR analysis of ecosystem productivity has also proven to be a reliable method for assessing mammalian habitat potential (Batchelor et al. 2023). Another essential means of measuring the impact of these droughts on ecosystems is to utilize remote sensing applications that measure biomass in relation to fluctuating temperatures (Cavanaugh et al. 2019). These techniques and technologies were not utilized in this study but would be valuable for future studies of environmental changes in cypress-tupelo swamps in relation to the potential for wildlife habitat.

## Conclusion

This study sought to understand the complex relationship between climate change-induced drought and cypress-tupelo swamp productivity in the NACP of the southeastern United States. MODIS data were used to extract NDVI, LAI, and EVI vegetation productivity from various points of cypress-tupelo swamps. The PDSI and SPI drought indices and CHIRPS precipitation data were also analyzed for areas of cypress-tupelo swamps, leading to the finding that drought change impacts the productivity of swamps near the coast. We determined that precipitation levels are decreasing throughout the southern range of the NACP and coastal cypress-tupelo swamp ecosystems and decreasing the net primary productivity of bald cypress and tupelo trees. Meanwhile, more inland and northern areas generally displayed increases in both wetness and vegetation productivity. An analysis of time series graphs and linear regressions for all swamp points determined these changes in swamp habitats across the NACP. Overall, our findings indicate that cypress-tupelo swamps in the south are more prone to drought, impacting their vegetation productivity and the overall health of these ecosystems. Future studies should further investigate the threat of drying climate conditions within the United States and their impacts on wetland ecosystems and the species that depend on them.

## Acknowledgments

This research was supported by the University of Southern Mississippi Gulf Coastal Plain Ecology Research Experience for Undergraduates which is funded by the National Science Foundation (2349627). We would also like to acknowledge Alyssa Merry for her feedback on the data analysis of this project.

