## Supplementary Information for "Drought and Productivity Trends in Freshwater Flooded Forest Swamps in the North American Coastal Plain"

SI *Figure 1: Decadal trends in NDVI, EVI, and LAI for the 2500 sample plots of the 500 largest NACP freshwater flooded forests.*

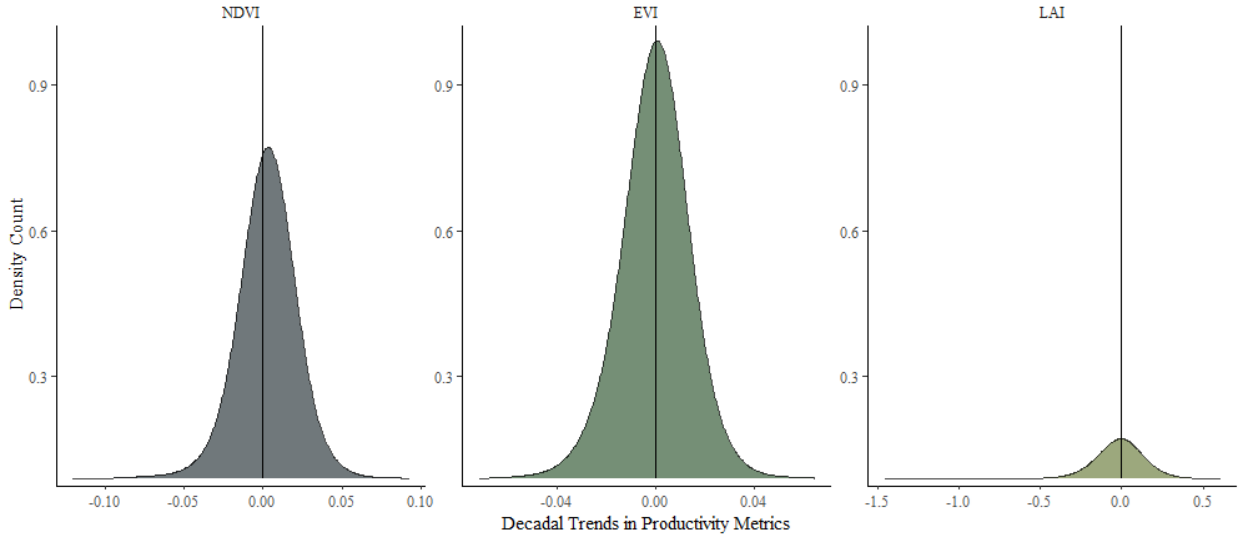

*SI Figure 2. Decadal trends in PDSI, Precipitation, and SPI based on annual trends from 2000 to 2024 for the 2500 sample plots of the 500 largest North American Coastal Plain freshwater flooded forests.*

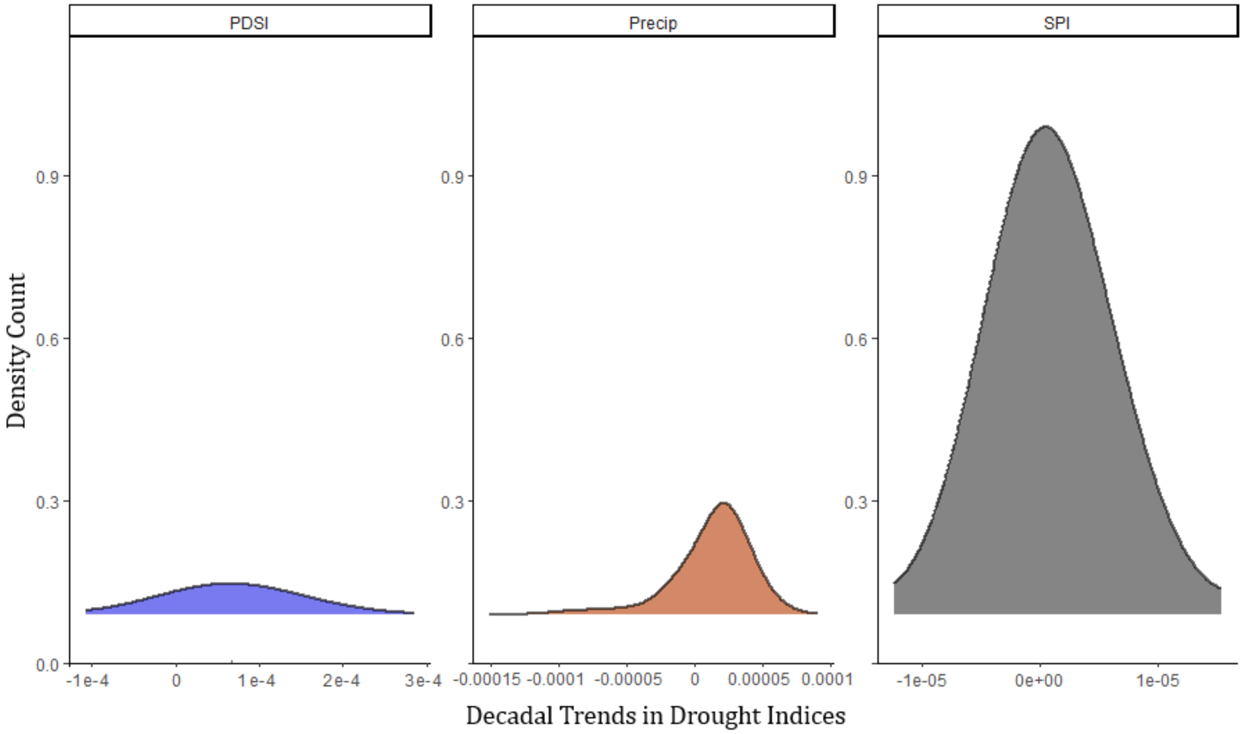

SI Figure 3: Productivity hotspots of a) precipitation and b) PDSI with red indicating statistical clusters of increasing precipitation and decreasing drought while blue indicates decreases in precipitation and increasing drought.

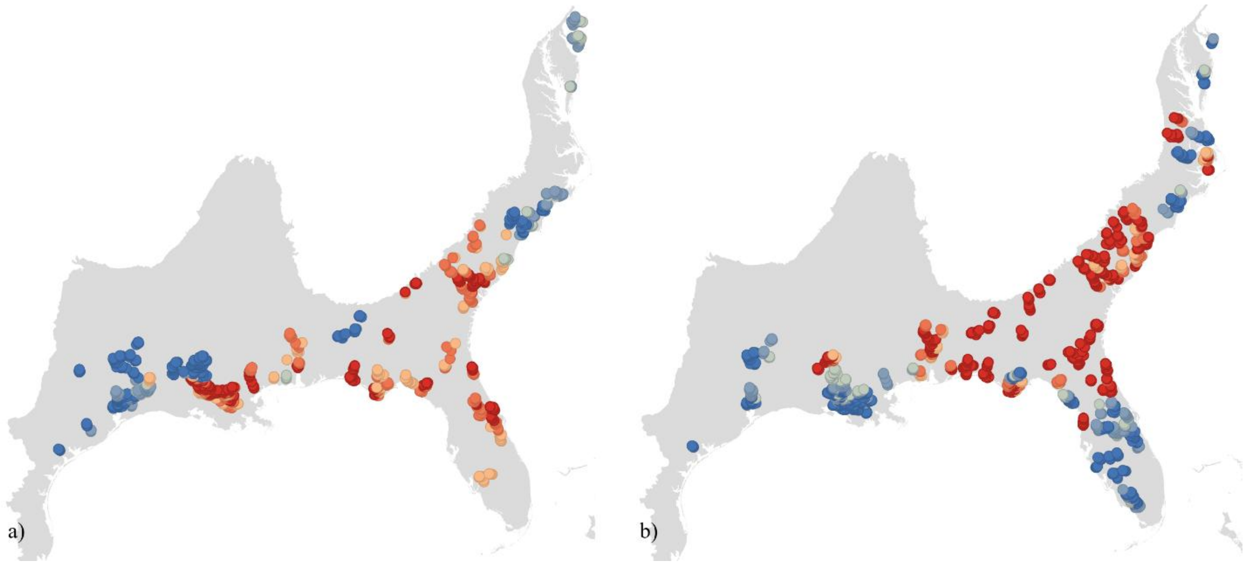

SI Figure 4: Reference map of the National Wildlife Management Areas and Refuges. Data taken from PAD-US.

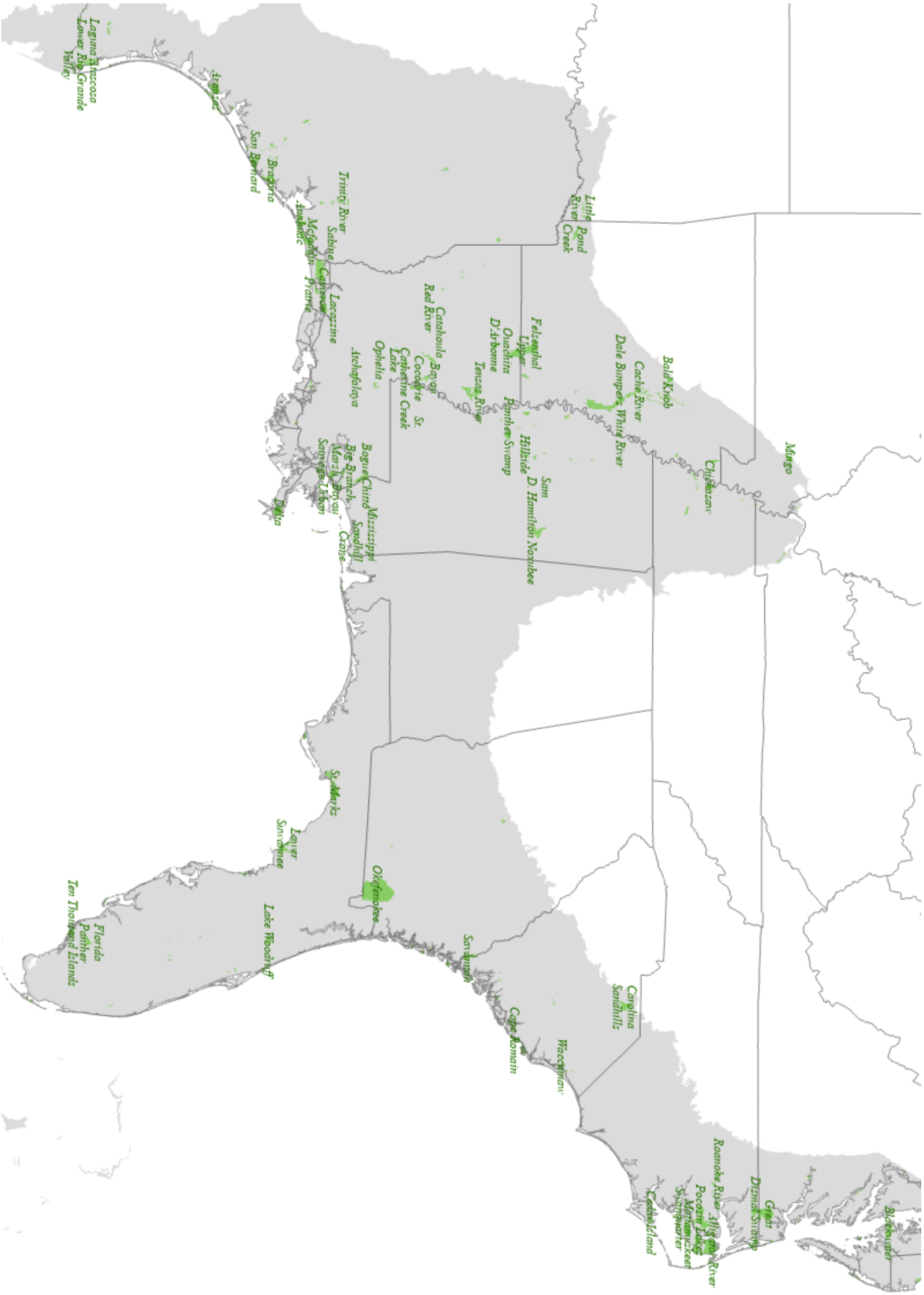

SI Figure 5: Reference map of the major national highways and interstates in the USA. Data taken from the Department of Transportation.

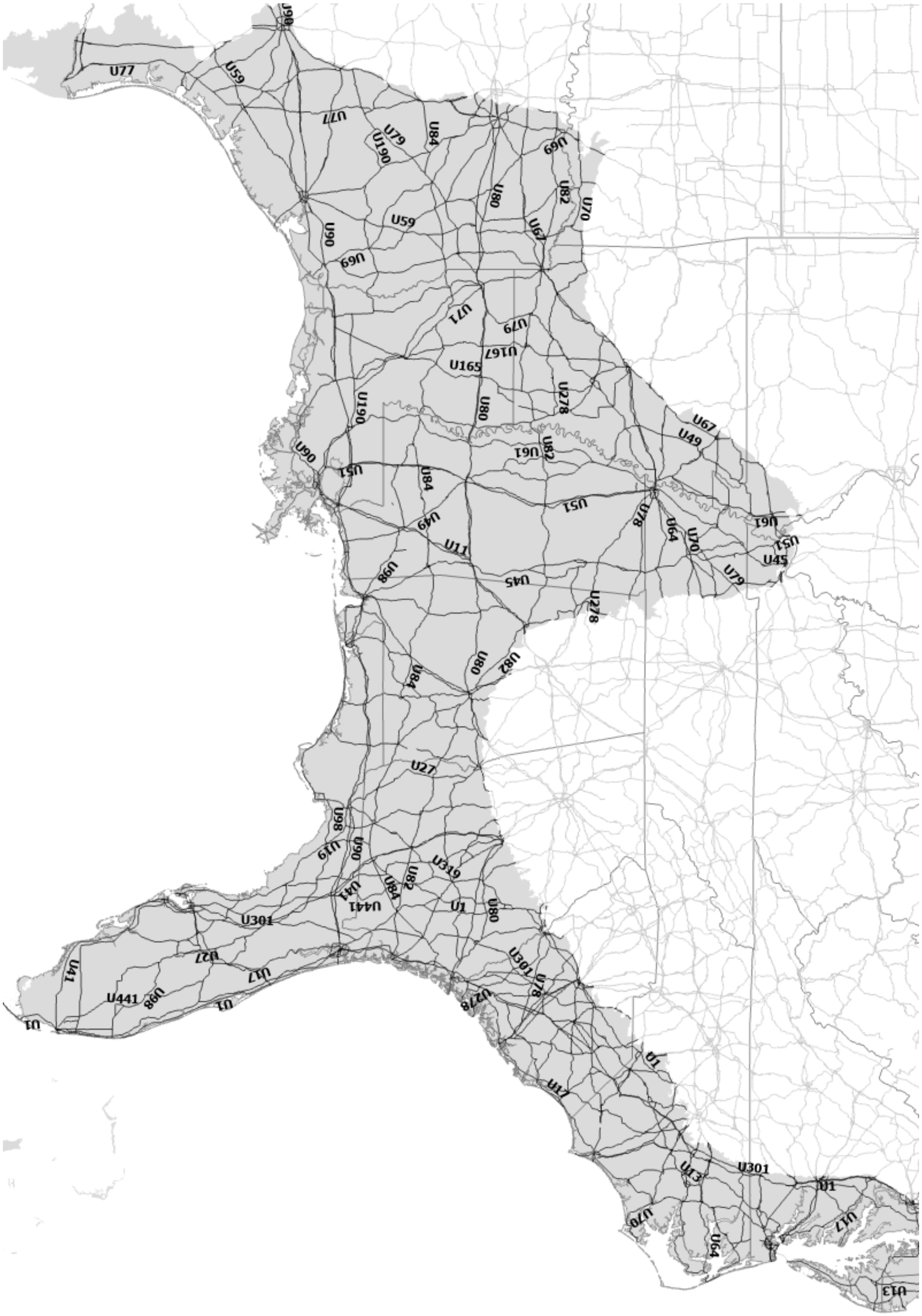

SI Figure 6: Reference map of the major rivers tributaries in the USA. Data taken from the Department of Transportation.

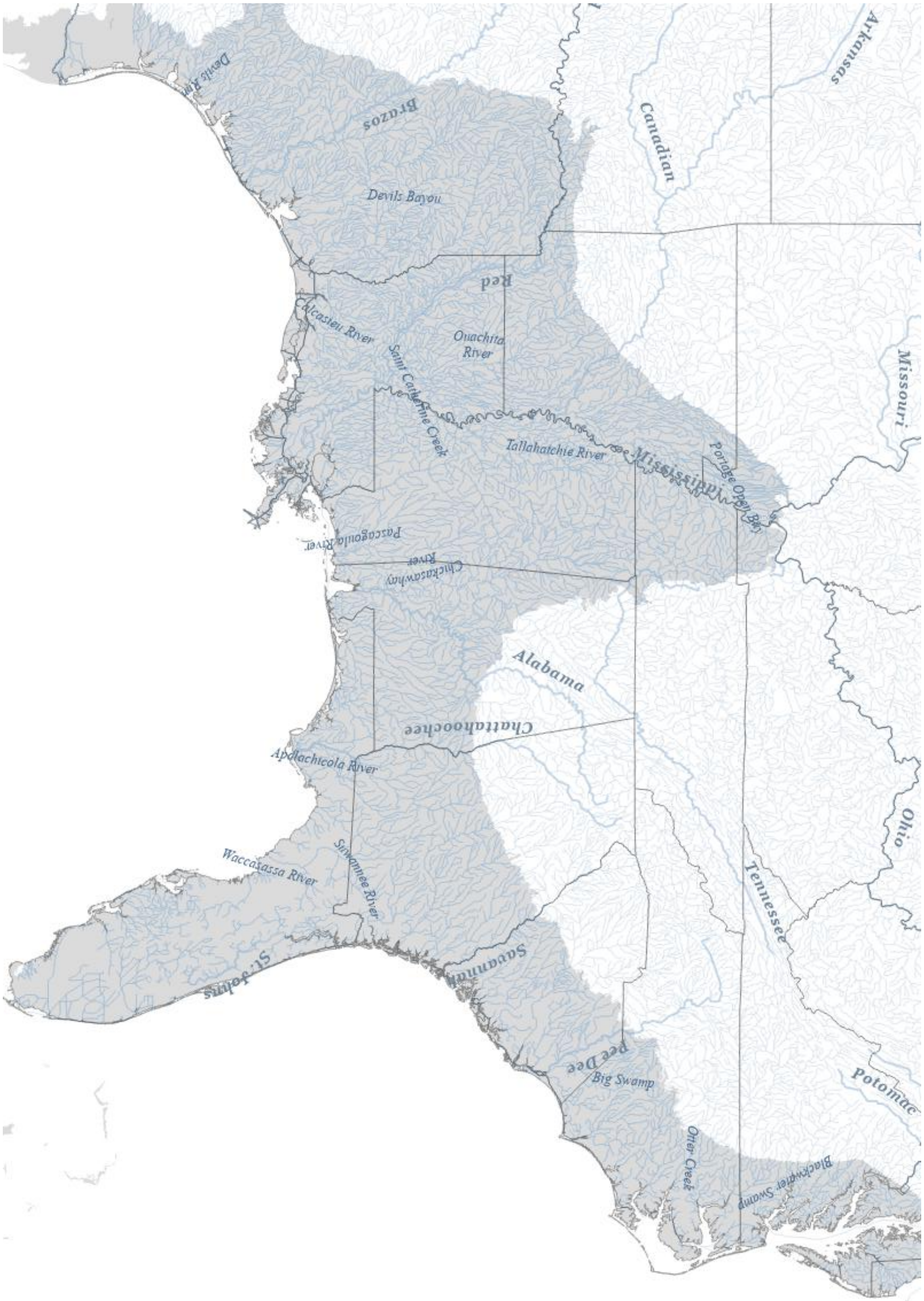

SI Figure 7: Reference map of the select cities in the North American Coastal Plain. Data compiled by ESRI based on data from the US Census Bureau, Department of Commerce, NOAA, and National Geodetic Survey.

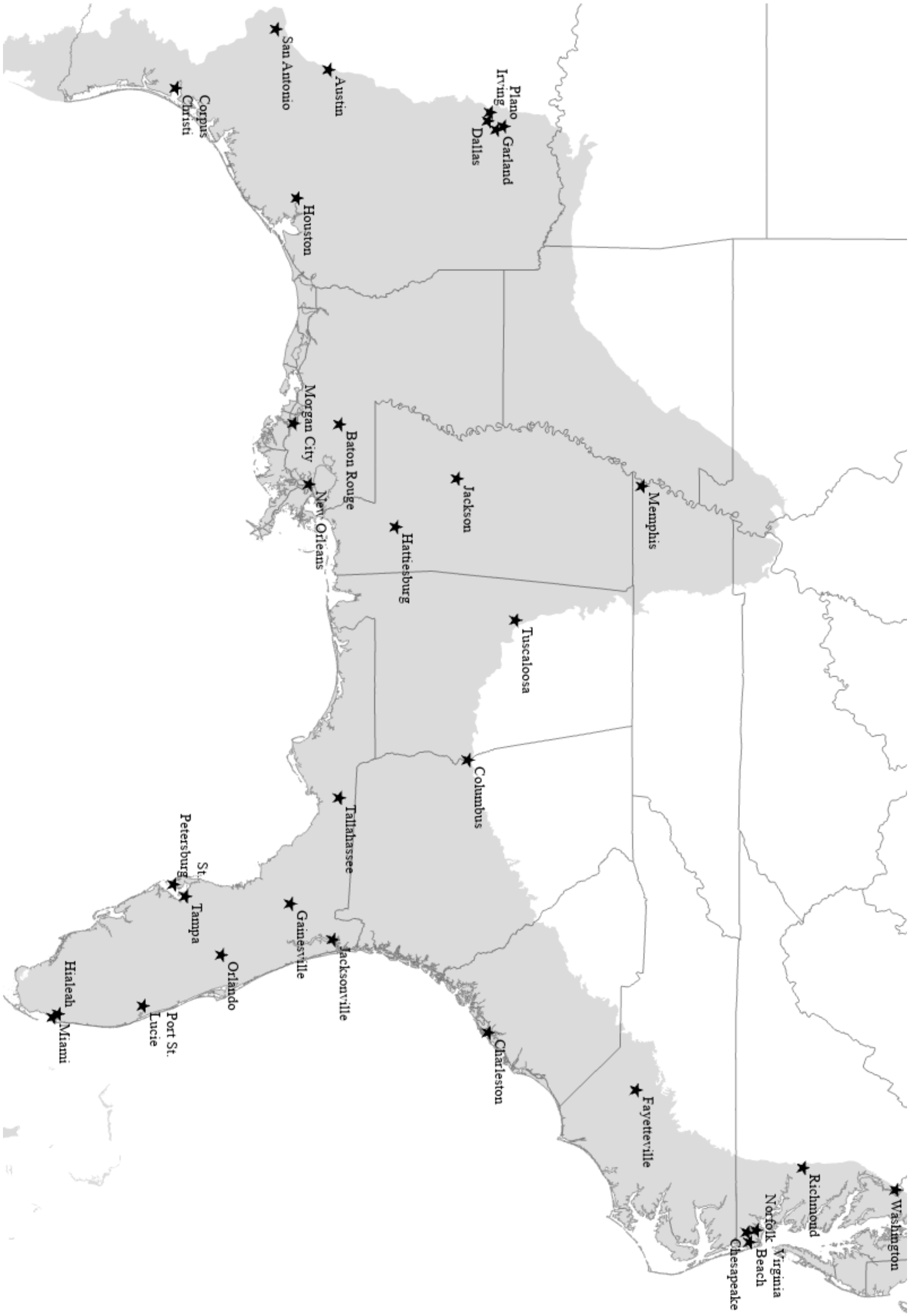
